# All-Atom Modeling and Simulation of Biopolymer Interface: Dual Role of Antifouling Polymer Brushes

**DOI:** 10.1101/2025.05.02.651893

**Authors:** Seonghan Kim, Zhefei Yang, Jacek Jakowski, P. Ganesh, Scott Retterer, Jan-Michael Carrillo

## Abstract

Antifouling polymers are highly valuable in a variety of applications, including antiviral coatings, targeted drug delivery, and marine coatings, where preventing unwanted protein adsorption is critical. Although extensive experimental studies have characterized polymer-protein interactions, computational studies remain limited due to the inherent difficulty of modeling and integrating these distinct components within a unified system. This study presents molecular modeling and simulation of polyelectrolyte and polyzwitterionic brushes—poly(dimethylaminoethyl methacrylate) (PDMAEMA), poly(2-(N-oxide-N,N-dimethylamino)ethyl methacrylate) (PNOMA), and poly(2-(N-3-sulfopropyl-N,N-dimethylammonium)ethyl methacrylate) (PSBMA)—grafted onto α-quartz substrates in the presence of lysozyme protein. The brush models were developed to closely replicate experimentally synthesized brush samples and to provide detailed insights into structural and dynamical changes at the molecular level during protein adsorption. Using steered molecular dynamics simulations, we show that the PSBMA brush, due to its high local density, exhibits the greatest resistance to protein insertion. C*α* root-mean-square deviation and interaction patterns analyses further reveal that PSBMA also induces the most significant destabilization of lysozyme, while PDMAEMA brush enhances protein stability through ion-mediated interactions. The PNOMA brush, while requiring the lowest force for protein adsorption, induces greater protein destabilization than the PDMAEMA brush primarily due to electrostatic repulsion caused by a short carbon spacer length. To the best of our knowledge, this study presents the first comprehensive and realistic model system—comprising nanomaterials, polymers, and proteins, specifically antifouling polyzwitterions and polyelectrolytes grafted onto *α*-quartz substrates via linkers. These findings highlight the dual role of antifouling polymer brushes: resisting protein adsorption and modulating protein structural dynamics, offering valuable insights for the rational design of next-generation antifouling materials.

**TOC GRAPHICS:** 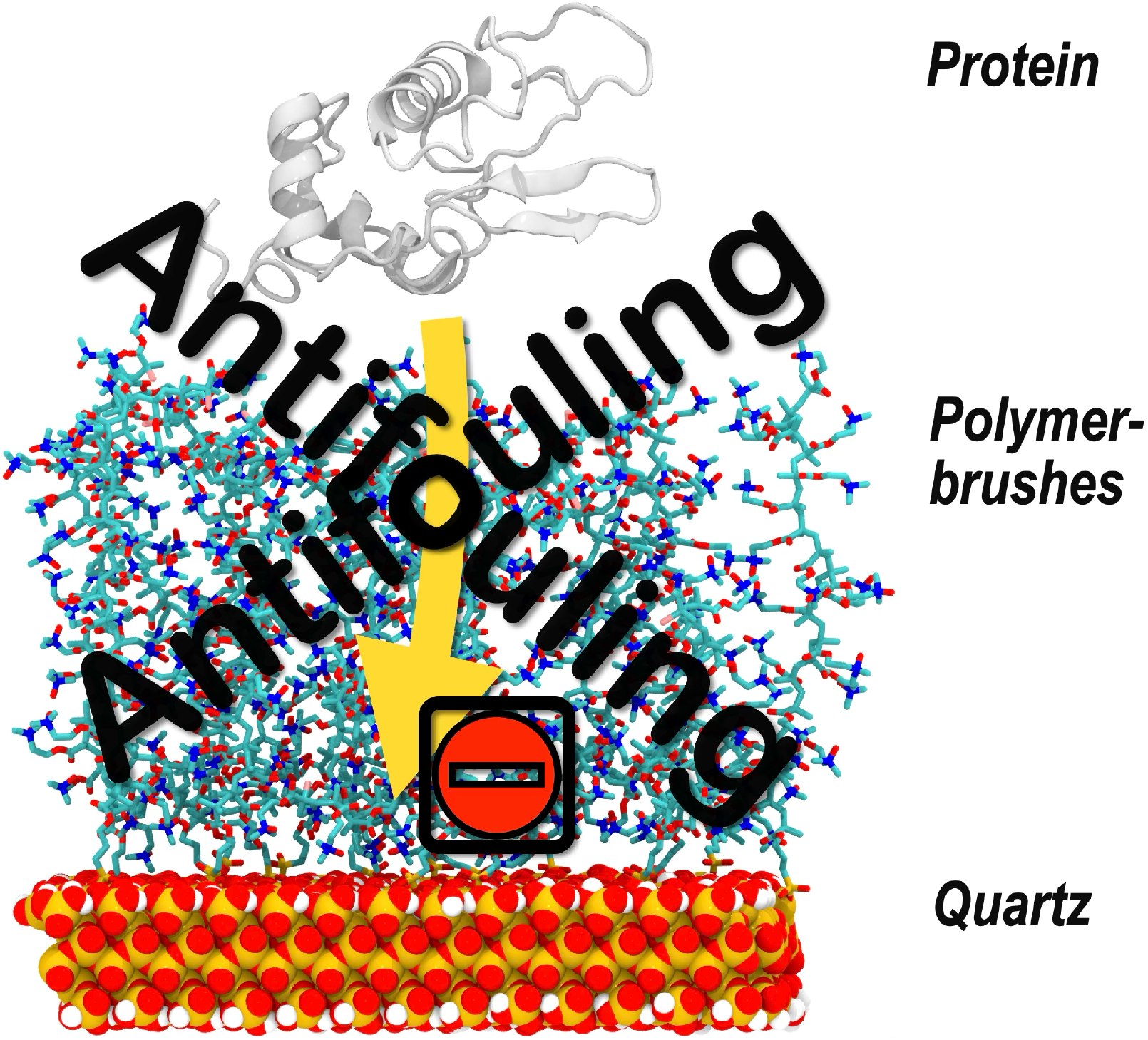

## 1. INTRODUCTION

Antifouling polymers are critical in a wide range of applications, including antibacterial coatings,^1,2^ nanocarriers for drug delivery,^3-5^ biosensors,^6,7^ marine antifouling coatings,^8,9^ and blood contacting devices.^10,11^ Preventing unwanted protein adsorption is essential for these applications, and polyzwitterions—polymers that contain both positively and negatively charged groups within their monomer units—are promising antifouling materials due to their charge balance, hydrophilicity, biocompatibility, and superior protein resistance.^4,12-14^ Over the past two decades, polyzwitterions have attracted considerable attention in antifouling applications due to their strong ionic solvation with water, in contrast to the weaker hydrogen bonding interactions of traditional poly(ethylene glycol) (PEG) or PEG-based materials, which have long been considered the gold standard in antifouling materials.^4^

Despite significant experimental progress, with over 800 SCI-indexed publications on polymer brushes published annually over the past decade,^15^ computational studies have remained limited, particularly those investigating interactions between proteins and polymer brushes. Fewer than 100 computational studies are published each year,^15^ likely due to challenges such as the need for interdisciplinary expertise, complexity of modeling multicomponent systems, the limited availability of suitable force fields, and differences in time and length scales of components involved. This disparity has led to various experimental observations remaining unexplained by current computational approaches, limiting our understanding of protein–polymer interactions and hindering the rational design of antifouling materials.

To address this gap, atomistic molecular dynamics (MD) simulations have emerged as a powerful tool for characterizing biomolecular interactions at atomistic resolution. While still somewhat limited in the context of protein dynamics within antifouling polymer brushes, MD simulations have been widely applied to study protein interactions with diverse components, including ligand,^16^ other proteins,^17-20^ nanomaterials,^21^ and polymers,^22,23^ which are often inaccessible through experimental techniques. By providing residue-specific insights, MD can significantly enhance our understanding of protein behavior at biopolymer interfaces and inform the rational design of antifouling materials, ultimately reducing reliance on costly and time-consuming experimental screening.

Although only a limited number of studies have applied MD to model polymer brush systems, several efforts have demonstrated its value at the molecular level. For example, Santos et al. reported the swelling and collapsing behavior of poly(dimethylaminoethyl methacrylate) (PDMAEMA) and poly(2-(methacryloyloxy)ethyl trimethylammonium chloride) (PMETAC) in response to changes in pH, solvent, and salt concentrations using atomistic MD simulations.^24^ Their study provided molecular-level insights into the conformational dynamics of the polymer brushes, although the model simplified the silicon substrate by representing it with sulfur atoms under positional restraints.

In 2019, Liu et al. reported the optimal packing structures of polymer brushes via unit cell approaches with a gold monolayer substrate.^25^ Using poly(carboxybetaine methacrylate) (PCBMA), poly(2-methacryloyloxyethyl phosphorylcholine) (PMPC), poly (2-(N-3-sulfopropyl-N,N-dimethylammonium)ethyl methacrylate)) (PSBMA), and PEG, they characterized brush structures and ranked antifouling performance by conducting steered molecular dynamics (SMD) simulations. However, the relatively short brush models used in their study, five repeating units, were insufficient to capture the full structural and dynamic behavior of the brushes during the protein adsorption, limiting conclusions to surface-level interactions.

Huang et al. employed trimethylamine N-oxide (TMAO) as a zwitterionic molecule and examined its interfacial behaviors, both experimentally through sum frequency generation (SFG) vibrational spectroscopy and computationally with MD simulations.^26^ They investigated surface hydration, salt effects, and protein interactions, confirming the role of carbon spacer length (CSL) in brush structures. Additionally, they monitored the dynamics of a lysozyme protein near the interface, observing the landing and rebounding motion of protein molecules as evidence of antifouling performance. An advancement in their study was the use of a more realistic *α*-cristobalite compared to earlier models. However, their polymer brush systems included a high fraction of CH_2_ alkyl spacers, which are not representative of typical antifouling polyzwitterions.

More recently, in 2024, Song et al. combined atomic force microscopy (AFM) and MD simulations to investigate salt-responsive antifouling behavior of PCBMA, PMPC, and PSBMA polyzwitterion brushes.^27^ The findings revealed the influence of ionic valance (e.g., Na^+^ vs. Ca^2+^) and ion concentration on surface potential, hydration behavior, and protein adhesion. While informative, their computational model was limited by the use of a single-layer silicon plate with fixed terminal carbon atoms, which lacked realistic surface heterogeneity and grafting geometry.

While these studies have contributed important insights into polymer brush swelling, packing, hydration behavior, and protein interactions, they often rely on simplified models—such as idealized substrate, restrained or fixed components, and short or simplified polymer chains—and are not integrated with detailed analyses of protein conformation. In particular, none of these studies has successfully captured the structural and dynamical changes of both polymer brushes and proteins within a realistic brush environment. This remains a critical gap in achieving a comprehensive understanding of brush-protein interactions, which is essential for rational design and applications of antifouling polymers.

To overcome these challenges, we developed, for the first time, a comprehensive and realistic all-atom model system consisting of nanomaterials, polymers, and proteins. Specifically, antifouling polyzwitterions and polyelectrolytes were grafted onto *α*-quartz substrates via linkers, closely replicating experimental conditions, and studied together in the presence of lysozyme protein. Through quantum calculations, MD, and SMD simulations, we characterized three representative polymer brushes, PDMAEMA, poly(2-(N-oxide-N,N-dimethylamino)ethyl methacrylate) (PNOMA), and PSBMA, to evaluate their antifouling performance against lysozyme adsorption. Furthermore, we analyzed protein stability and its interaction patterns within the brush environment, providing valuable atomistic insights into the role of antifouling polymer brushes in modulating both protein adsorption and stability.

## 2. METHODS

### 2.1. Modeling of Biopolymer Interface

Three polymer brushes were modeled using CHARMM^28^ script to investigate their antifouling properties, i.e., PDMAEMA, PNOMA and PSBMA (**Figure 1A–C**).

**Figure 1.**
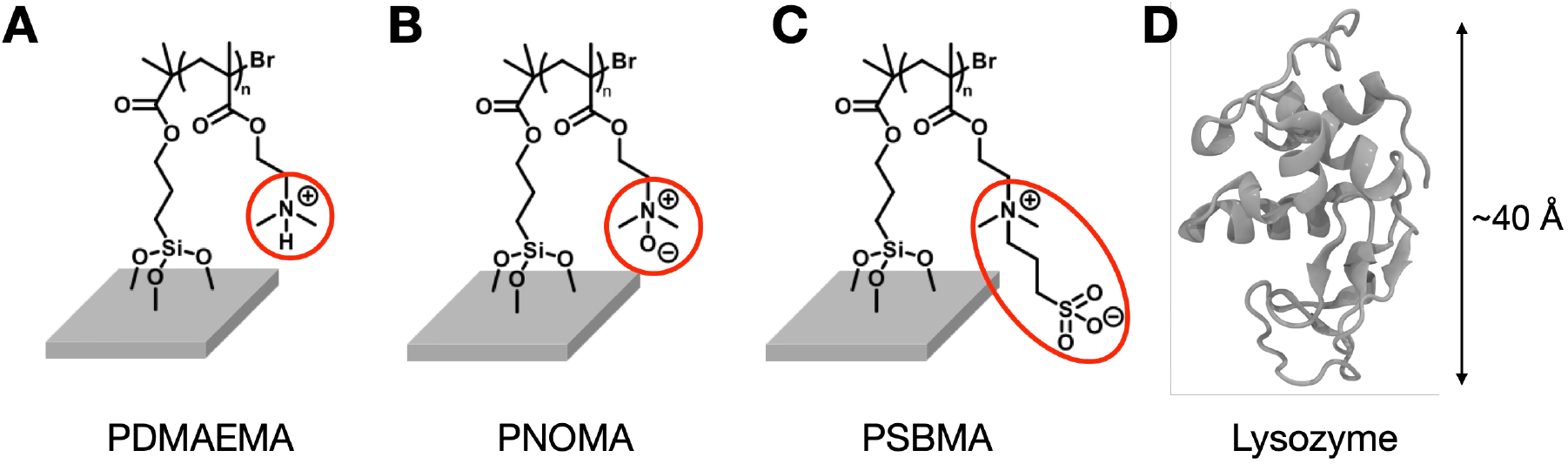
(A–C) Model brush structures and (D) a lysozyme (PDB ID: 7LYZ) used in this study at pH 7; (A) PDMAEMA, (B) PNOMA, and (C) PSBMA. The degree of polymerization, *n*, was set to 20. Structural differences between the three polymer brushes are highlighted with red circles.

#### 2.1.1. Polymer Brush

*α*-Quartz, the most stable and prevalent form of a silica and abundant in nature, is widely utilized in applications such as electronics, optics, and industrial processes.^29-31^ In this study, *α*-quartz was used as a substrate for brush structures to mimic the experimental condition. The *α*-quartz surface was constructed with (001) surfaces oriented along the Z-direction using the CHARMM-GUI *Nanomaterial Modeler*.^21,32^ For model parameters such as bonded and nonbonded parameters, the Interface Force Field (IFF),^33^ which was developed for simulations of biotic–abiotic interfaces, was selected due to its accuracy and compatibility with other force fields (FFs) used in this study. The model dimensions were set to 63.9 × 68.1 × 10.9 Å^3^ with 13% of surface ionization (equivalent to pH 7), which involves surface modification of oxygen atoms into silanol groups. Following model construction, 150 ns of MD simulations were performed for equilibration, and the final simulation trajectory was taken to use the substrate as an initial model for building the polymer brush.

After the substrate was modeled, the monomers of PDMAEMA, PNOMA, and PSBMA, as well as the linker connecting the polymer brushes to the surface were constructed. Bond formation was implemented using the PATCH option in CHARMM. Parameters for these structures were adapted from CGenFF and organosilane FF,^34-37^ as their chemical structures have been extensively validated for similar compounds. The degree of polymerization, *n*, was set to 20 to achieve the desired polymer length, which allows lysozyme to be fully inserted into the polymer brush during the SMD simulation, capturing entire interactions between the protein and the brushes (**Figure S2**).

Grafting density is one of the important parameters that determines the antifouling performance of polymer brushes. In this work, a grafting density of 0.4 chains/nm^2^ was selected, mimicking high grafting densities from experimental conditions.^38^ To achieve this density, 17 polymer chains, each consisting of a polymer and a linker, were randomly placed on the silica surface. Physical bonds were constructed using the PATCH command in CHARMM.^28^ After building all polymer brush models (PDMAEMA, PNOMA, and PSBMA), at least 200 ns of MD simulations were performed for each system equilibration, with no changes in brush thickness were observed after 100 ns. The final simulation trajectories were used as initial brush structures for SMD simulations.

#### 2.1.2. Lysozyme Protein

Lysozyme is a ubiquitous protein found in many biological fluids, characterized by its relatively smaller size (14.3 kDa with 129 amino acid residues, **Figure 1D**). Due to these features, it is frequently used as a model protein in many studies.^25,26,39-41^ In this work, a lysozyme (PDB ID: 7LYZ) with net charge of +8e at pH 7 was used to test antifouling properties of the polymer brushes.

To equilibrate the lysozyme prior to conducting SMD, a solvated system containing lysozyme, ions, and water was constructed using CHARMM-GUI *Solution Builder*.^32^ Once the model was built, MD simulations were performed for 200 ns for its equilibration with CHARMM36(m) FF.^42,43^ The simulation was monitored by calculating C*α* root-mean-square-deviation (RMSD) tracking stability of the protein, and the final simulation trajectory was employed as the initial lysozyme model for incorporation into SMD systems.

#### 2.1.3. System Assembly

Once each component was equilibrated, all components were assembled into a single system using a CHARMM script. The initial distance between the centers of masses (COMs) of the lysozyme and the *α*-quartz was carefully adjusted to ensure that the protein did not touch the brush interface prior to the SMD simulation. The assembled system was then solvated with water containing 150 mM KCl to mimic physiological condition using CHARMM-GUI *Multicomponent Assembler* and *Input Generator*.^23,44^

### 2.2. Multi-Scale Simulation

#### 2.2.1. Quantum Mechanical Calculations

For the PNOMA model, quantum mechanical simulations were performed due to its unique structure, which includes an amine oxide—a moiety that is uncommon in biological systems and thus not well-represented by general force fields such as CGenFF (**Figure 1B**). ^34-36^ Calculations were conducted using NWChem software,^45^ and the partial charges were obtained using the restrained electrostatic potential (RESP) method^46^ after geometry optimization. The B3LYP functional with the 6-311++G(d,p) basis set was employed for these calculations (**Figure S1B**).^47,48^ A monomer structure was used in the simulation (**Figure S1A**), and partial charges around the backbone atoms were adjusted to neutralize the polymer. To validate these results, the radial distribution function, g(r), between the oxygen of water and the oxygen and nitrogen atoms of the polymer was calculated, showing consistency with previously reported studies (**Figure S1C**).^49^

#### 2.2.2. Molecular Dynamics Simulation

MD simulations were conducted using NAMD.^50^ The CGenFF,^34-36^ IFF,^33^ and CHARMM36(m) ^42,43^ FFs were used for polymer, silica, and lysozyme, respectively, with the TIP3P water model.^51^ The van der Waals interactions were handled with a force-based switching function, smoothly tapered from a 10 to 12 Å cutoff.^52^ Electrostatic interactions were computed using the particle-mesh Ewald (PME) method^53^ with a 1 Å grid spacing with a sixth order b-spline. Bond lengths involving hydrogen atoms were constrained using the SHAKE algorithm.^54^

The NVT ensemble with positional and dihedral restraints were applied and gradually decreased for the equilibration processes. Then, the production run was conducted in the NPT ensemble with periodic boundary conditions at 303.15 K and 1.01325 bar, controlled via Langevin dynamics (damping coefficient of 1 ps^-1^) and Langevin piston method, respectively.^55^

#### 2.2.3. Steered Molecular Dynamics Simulation

NAMD with the COLVARS method^50^ was used for SMD simulation. The lysozyme was initially positioned above the polymer brush and aligned along the Z-axis. A harmonic pulling force was applied along the Z-direction to the COM of the protein’s Cα atoms to guide its adsorption toward the quartz surface. This setup allowed the protein to translate along the Z-axis while remaining free to rotate in the X and Y directions. The effective pulling force was computed using the equation: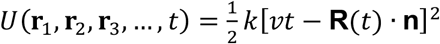, where *k* is the spring constant, *v* is the moving speed of the dummy atoms (i.e., spring potentials), **R**(*t*) is the position of the COM of the lysozyme, and ***n*** is the unit vector defining the pulling direction along Z. This force allows the spring-connected proteins to be pulled into the brush surface. A pulling speed of 0.5 Å/ns and a spring constant of 15 kcal/mol/Å^2^ were used to ensure gentle yet effective sampling. To improve statistical results, five independent simulations were performed for at least 150 ns, ensuring full insertion of the lysozyme into the polymer brush.

## 3. RESULTS AND DISCUSSION

### 3.1. Force Profiles During the Protein Adsorption onto Polymer Brushes

SMD simulations were conducted to investigate antifouling properties of PDMAEMA, PNOMA, and PSBMA brushes. During the simulations, a lysozyme was pulled toward, and into, the polymer brushes, with the hypothesis that stronger antifouling brushes would require greater forces for protein insertion (**Figure 2A**). Initially, the lysozyme was located on top of brushes, where its initial COM-COM distance, *D*, was set to 60 Å. As the simulation progresses, the protein contacts the brush interface around *D*=55 Å (**Table 1**). Once it passes that point, it begins to penetrate the interface. The results show that PSBMA requires the highest forces for protein insertion, followed by PDMAEMA and PNOMA brushes. Stronger pulling forces indicate that there are more repulsive interactions between the lysozyme and polymer brushes, which suggests stronger antifouling properties.

To understand the differences in the force profiles, a density analysis was performed (**Figure 2B**). The results show that the PSBMA brush has the highest density, making it the bulkiest among the three brushes (see also **Figure S3**). Interestingly, while the PNOMA brush exhibits a higher density than PDMAEMA, it has a thinner brush thickness, despite the only difference being an O atom in PNOMA and H atom in PDMAEMA, both attached to the N atom.

**Table 1.**
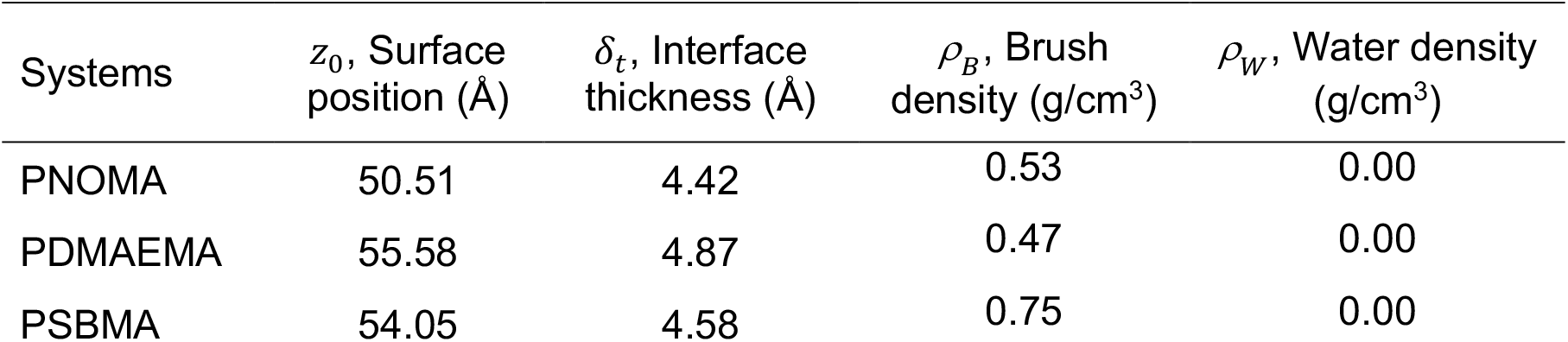
Surface position (*z*_0_) and interface thickness (*δ*_*t*_) of each polymer brush fitted from the hyperbolic tangent fit.

| Systems | $z_0$, Surface position (Å) | $\delta_t$, Interface thickness (Å) | $\rho_B$, Brush density (g/cm<sup>3</sup>) | $\rho_W$, Water density (g/cm<sup>3</sup>) |
| --- | --- | --- | --- | --- |
| PNOMA | 50.51 | 4.42 | 0.53 | 0.00 |
| PDMAEMA | 55.58 | 4.87 | 0.47 | 0.00 |
| PSBMA | 54.05 | 4.58 | 0.75 | 0.00 |

**Figure 2.**
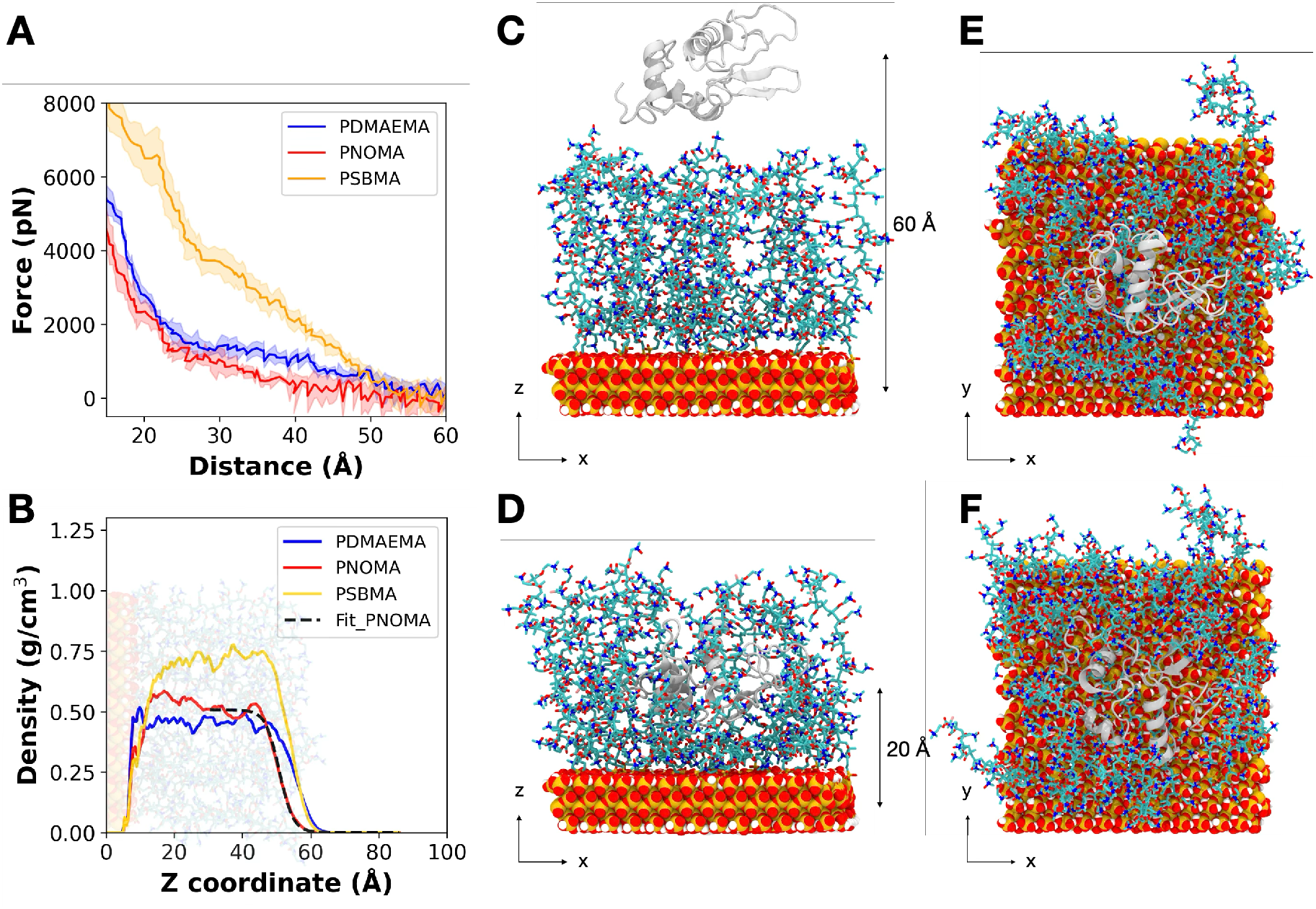
(A) Force profiles during the SMD simulation, representing the protein adsorption process (i.e., from *D*=60 Å to 20 Å) on each polymer brush (red: PNOMA; blue: PDMAEMA; yellow: PSBMA). (B) Density profiles of polymer brushes for each system, with the fit of PNOMA brush shown as black dashed lines. Fits for PDMAEMA and PSBMA are omitted for clarity. (C–F) Representative snapshots at *D*=60 Å and *D*=20 Å of the PNOMA brush. The distance, *D*, refers to COM separation between lysozyme (light gray) and the quartz substrate (orange: Si; red: O; white: H). Side views (C,D) and top views (E,F) are shown, where (C,E) corresponds to the initial state (*D*=60 Å) and (D,F) to the final state (*D*=20Å). H atoms in the brush were hidden for clarity. See **Figure S2** for additional snapshots captured at 10 Å intervals.

For a further quantification, brush properties, such as surface position and interface width, were calculated by fitting a hyperbolic tangent function to the density profiles of each polymer brush (without a lysozyme):^56-58^

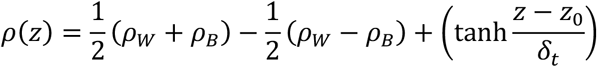

where *ρ*_*W*_ and *ρ*_*B*_ are the densities of water and polymer brushes, and *z*_0_ and *δ*_*t*_ represent the surface position and interface thickness. The fitting results are summarized in **Table 1**, and the fit of PNOMA brush is shown in **Figure 2B** as dashed lines. For clarity, the fits of PDMAEMA and PSBMA are not shown. From the fit data, we confirm that the PDMAEMA brush has the highest brush thickness (surface position) and interface width (interface thickness), followed by PSBMA and PNOMA brushes.

### 3.2. 2D Density Map of Polymer Brushes Before and After Protein Adsorption

To better understand changes in brush behavior after protein adsorption, 2D density maps were compared before (**Figure 4A–C**) and after the adsorption (**Figure 4D–F**) for each polymer brush; a clear difference was observed between polyelectrolytes (PDMAEMA) and polyzwitterions (PNOMA and PSBMA).

**Figure 4.**
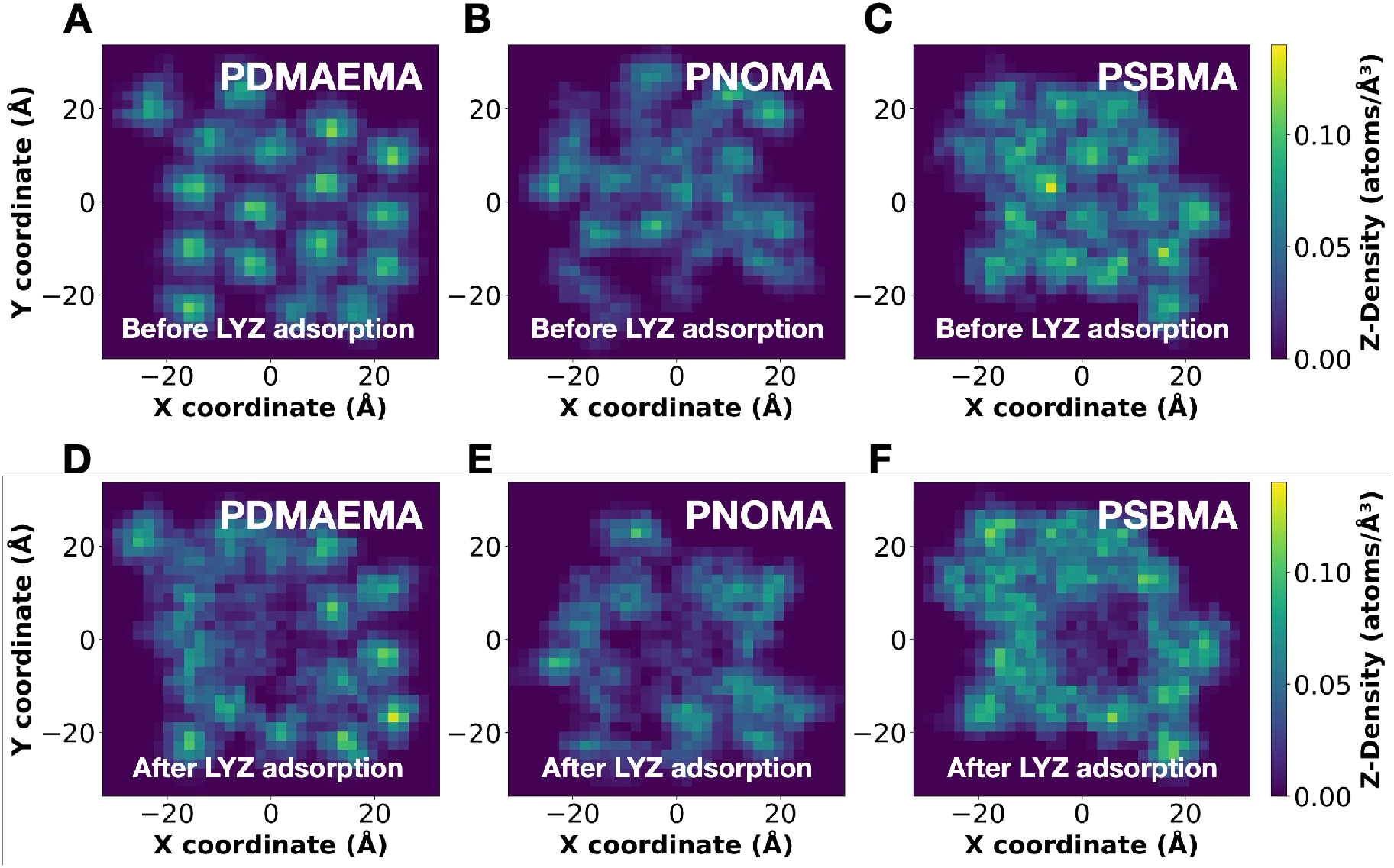
2D brush density maps of (A–C) before and (D–F) after lysozyme adsorption on (A,D) PDMAEMA, (B,E) PNOMA, and (C,F) PSBMA brushes. The X- and Y-axes correspond to lateral coordinates across the substrate surface. Lighter Z-density colors indicate higher density, while darker colors indicate lower density at each coordinate.

In the PDMAEMA brush, an empty space between each chain is observed (**Figure 4A**). This gap can be attributed to having positive charges at N atoms (**Figure 1A**), causing them to extend in the Z-direction by elongating their backbone through electrostatic repulsive interactions. This extension is found by relatively high Z-density observed across the brush (brighter colors indicate higher Z-density), further confirming the thickest surface position and interface thickness of the PDMAEMA brush (**Table 1**).

Compared to PDMAEMA brushes, the zwitterionic PSBMA brush exhibits wide-spanning distribution along the X- and Y-coordinates (**Figure 4C**). This is particularly due to its longer monomer CSL, defined by the number of carbon atoms between the positive and negative charged atoms in the monomer (CSLs of PSBMA and PDMAEMA are 3 and 0, respectively; see **Figure 1**). A longer CSL increases bulkiness within the brush (**Figure 2B**,**S3**) ultimately increasing antifouling performance of the polymer brush as reported by previous studies.^12,59,60^ As a result, the PSBMA brush requires the highest forces for lysozyme penetration during the course of SMD simulation. It is worthwhile to mention that a recent machine learning (ML) study trained on experimental data reported that the grafting density and molecular weight have the greatest impact on the antifouling properties among 12 features, including pH, ionic strength, flow rate, and density, with density having the strongest effect.^38^ Notably, with the same degree of polymerization, the PSBMA also possesses the highest density (**Figure 2B**), as well as molecular weight among the three polymers (monomer molecular weights for PDMAEMA, PNOMA, and PSBMA: 158.22, 173.21, and 279.36 g/mol, respectively). While our definition of density refers to effective local brush packing, within the same grafting density of 0.4 chains/nm^2^ across all systems, this interpretation aligns with the ML findings, further supporting that the PSBMA is likely to achieve superior antifouling performance due to higher local density.

The PNOMA brush, a polyzwitterion, also spans widely across the silica substrate without showing vacant space between chains like the PDMAEMA (**Figure 4B**). This is due to the PNOMA brushes’ ability to interact with each other through their charge–charge interactions, making them more flexible on the surface and thus reducing its thickness. This is also validated by its density profile, a denser but thinner brush than PDMAEMA, (**Figure 2B**) and the fit data (**Table 1**), smaller surface position.

As noted earlier, the PDMAEMA brush exhibits the second-highest force profiles during the SMD simulations. Under these simulation conditions mimicking the experimental condition of pH 7, a lysozyme contains +8e charges. This charge distribution can negatively impact the protein penetration through repulsive interactions between the positively charged lysozyme and positive amine groups in the polycationic PDMAEMA brush. Thus, a comparison of the 2D density maps before and after protein adsorption for the PNOMA (**Figure 4B,E**) and PDMAEMA (**Figure 4A,D**) brushes reveals that the PDMAEMA brush shifts laterally in the X-Y plane due to electrostatic repulsion from the positively charged lysozyme. In contrast, the PNOMA brush exhibits minimal lateral shifts, further confirming the repulsive interactions between lysozyme and the PDMAEMA brush responsible for showing higher force profiles than the PNOMA brush.

### 3.3. Influence of Polymer Brushes on Protein stability and Interaction Patterns

Characterizing antifouling polymers is important, but equally crucial is understanding how these polymers affect protein structure during adsorption and penetration. For instance, when developing antiviral surfaces by utilizing antifouling properties, a desirable outcome can be achieved if the protein loses its function through structural disruption. In such cases, a coating that induces structural changes on proteins can be an effective antiviral surface.

To evaluate protein stability after the adsorption, the C*α* RMSD was calculated as a function of COM distance between the protein and the silica substrate (**Figure 5A**). The RMSD results indicate that lysozyme exhibits the greatest structural changes in the PSBMA brush (RMSD>4Å), followed by the PNOMA, and PDMAEMA brushes. Interestingly, this trend does not align with the force curves shown in **Figure 2A**, where the order was observed to be: PSBMA > PDMAEMA> PNOMA. To further investigate this discrepancy between the PNOMA and PDMAEMA systems, we analyzed protein interaction patterns (**Figure 5B**).

**Figure 5.**
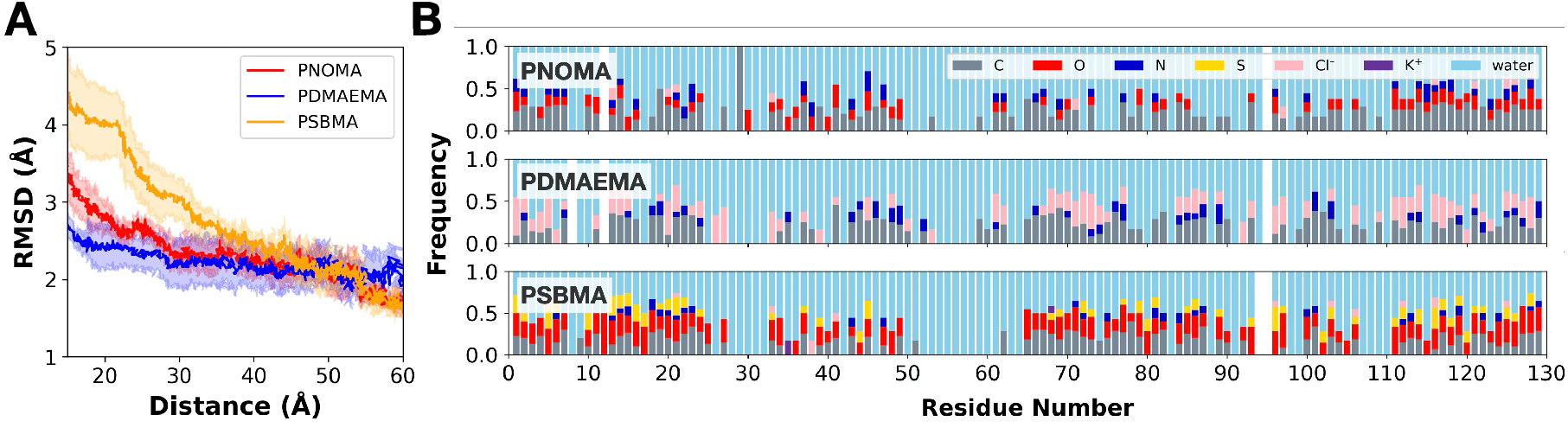
(A) The C*α* RMSD of a lysozyme in PNOMA (red), PDMAEMA (blue), PSBMA (yellow) brushes as a function of distance, *D*, between COMs of the lysozyme and the quartz. (B) The protein interaction patterns at *D*=20 Å when the protein is fully inserted into each brush (gray: C; red: O; blue: N; yellow: S; pink: Cl^-^; purple: K^+^; skyblue: water). Close contacts are defined as interactions within 4 Å between protein heavy atoms and the atoms of the polymer side chains, ions, and water molecules.

The interaction patterns can be investigated by calculating contact frequencies, defined as interactions occurring within 4 Å between protein heavy atoms and atoms in the side chains of the polymer brushes, surrounding ions, and water molecules. Specifically, for each heavy atom of the protein residues, pairwise distances were calculated with atoms such as C, N, O, and S from the brush (highlighted in **Figure 1**), along with nearby ions and water molecules. Then, the frequencies of these contacts were recorded for each residue.

This analysis was conducted at *D*=20 Å, where the lysozyme was fully embedded within the polymer brush (**Figure 2D,F**). In the PNOMA brush environment, lysozyme residues mainly interact with water, C, O, and N atoms in the brush (**Figure 5B**, top). Residues 12 and 95 show no such interactions, as they are oriented toward the hydrophobic inner region of the protein. The lysozyme, positively charged, frequently interacts with negatively charged O atoms via charge–charge interactions. However, a comparable number of interactions is also observed between the lysozyme and positively charged N atoms (CSL=0). While the O atoms contribute some stabilizing effects, the stability is largely negated by repulsive interactions with the neighboring N atoms.

In contrast, the PDMAEMA brush contains only positively charged N atoms, and it requires additional 340 Cl^-^ atoms for system neutralization (20 monomers per chain, 17 chains at 0.4 chains/nm^2^; see also **Figure S4** for ion distribution within the PDMAEMA brush). These excessive chloride ions form favorable interactions with the positively charged lysozyme, providing additional stabilization compared to the PNOMA brush (**Figure 5B**, middle). This enhanced stabilization reduces structural fluctuations in the lysozyme, as reflected by the lower RMSD values observed in the PDMAEMA brush system compared to the PNOMA brush (**Figure 5A**).

As demonstrated in the density analysis, the PSBMA brush exhibits a higher density and bulkier structure compared to the PNOMA and PDMAEMA brushes. As shown in **Figure 5A**, due to this crowded environment, lysozyme undergoes the most fluctuations during the initial stages of brush entry (*D<*40 Å), yet no significant structural changes that would be associated with protein unfolding were observed. This steric hindrance within the dense PSBMA brush is further supported by the interaction patterns (**Figure 5B**, bottom). A greater number of lysozyme residues exhibit additional contacts with the brush, and the contact frequencies are higher than those in the PNOMA and PDMAEMA systems. These results indicate that the increased local density of the PSBMA brush leads to more extensive interactions and heightened protein fluctuations, which could contribute to its superior antifouling performance compared to PNOMA and PDMAEMA brushes.

It should be noted that this observation could change under different environmental settings. For example, under biological conditions, typical ion concentrations range from 100 to 200 mM. However, the ion concentration used for developing lubricants and water-dispersible or aggregation-resistant materials can vary significantly, from 0 to 3000 mM.^61,62^ The charge of proteins is another factor to consider. If a negatively charged protein is tested in the PDMAEMA brush environment, Na^+^ ions (used to maintain the desired ionic strength) as well as the positively charged PDMAEMA brush itself will differently influence protein stability. The PSBMA brush interacting with the negatively charged protein might show even higher values for both force profiles and RMSD, due to negative charges in the SO_3_^-^ groups, which are separated by a relatively long CSL of 3. However, the effect of charge is likely less significant compared to density or ionic concentration, as reported by Okuyama et al.^38^ Therefore, careful approach is required when developing and testing antifouling performance, and these considerations remain to be investigated in future studies.

## 4. CONCLUSIONS

This study investigates the interactions between antifouling polymers and lysozyme protein. To the best of our knowledge, this represents the first comprehensive and realistic model system—comprising nanomaterials, polymers, and proteins, specifically antifouling polyzwitterions and polyelectrolytes grafted onto *α*-quartz substrates via linkers, along with a lysozyme protein—to be developed that closely replicates experimental conditions.

Using a multi-scale simulation approach that integrates quantum calculations, all-atom MD, and SMD, we characterized the structural and interfacial properties of three polymer brushes: PNOMA, PDMAEMA, and PSBMA. Among them, the PSBMA brush exhibited the highest local density, resulting in the greatest resistance to protein penetration. The PDMAEMA brush showed the largest brush height due to intra-brush electrostatic repulsion, while the PNOMA brush demonstrated more compact and laterally flexible conformation.

Protein stability upon adsorption within brushes was evaluated through Cα RMSD analysis and protein interaction patterns. The PSBMA brush, while requiring the highest forces for the protein adsorption due to steric hindrance, also caused the greatest structural fluctuations in lysozyme. Interestingly, despite requiring higher force for lysozyme adsorption, the PDMAEMA brush stabilized lysozyme more effectively than PNOMA, owing to favorable electrostatic interactions with excessive chloride counterions. These results suggest that the effects of charges are relatively small, yet not negligible on antifouling properties.

Overall, by introducing a realistic model system, this study provides new insights into the dual role of antifouling polymer brushes: resisting protein adsorption and modulating protein stability. These results suggest that the increased density of the PSBMA brush leads to more extensive interactions and greater protein fluctuations, ultimately contributing to its superior antifouling performance.

## ACKNOWLEDGEMENTS

This work was supported by the U.S. Department of Energy Office of Science FWP ERKCZ64, Structure Guided Design of Materials to Optimize the Abiotic-Biotic Material Interface, as part of the Biopreparedness Research Virtual Environment (BRaVE) initiative.

This work was performed at the Center for Nanophase Materials Sciences, the U.S. Department of Energy Office of Science User Facility operated at Oak Ridge National Laboratory. This research used resources of the Oak Ridge Leadership Computing Facility (OLCF) at Oak Ridge National Laboratory, which is supported by the Office of Science of the U.S. Department of Energy under Contract No. DE-AC05-00OR22725.

