## Supplementary information for "All-Atom Modeling and Simulation of Biopolymer Interface: Dual Role of Antifouling Polymer Brushes"

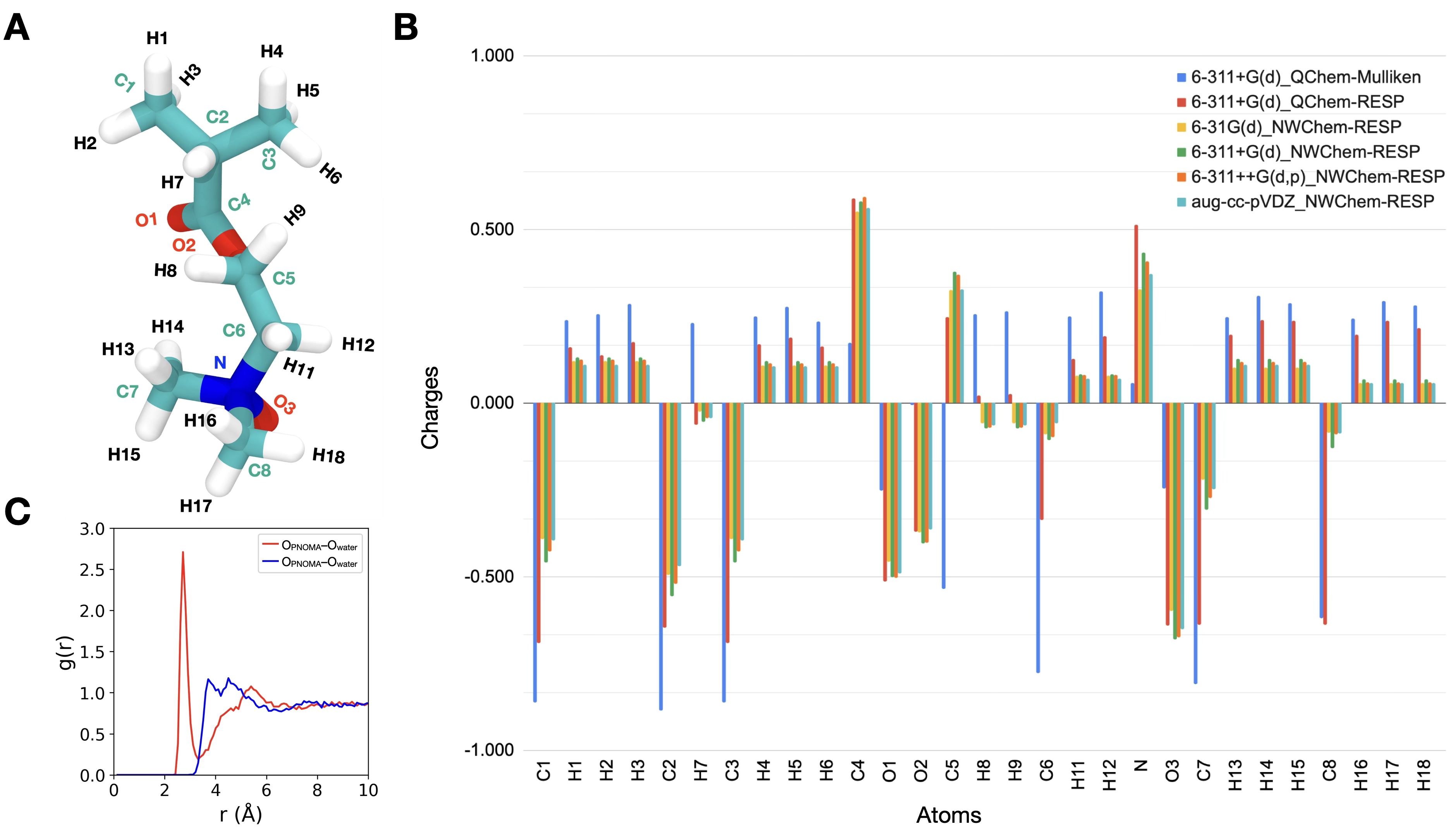

**Figure S1.** (A) An optimized monomer structure of PNOMA from DFT calculations using the B3LYP functional and the 6-311++G(d,p) basis set.^1,2^ (B) Benchmark results comparing various basis sets with Mulliken^3^ and RESP^4^ charge fitting methods, obtained from QChem^5^ and NWChem^6^ software. Partial charges corresponding to all atoms are shown in **Table S1**. (C) Radial distribution function, g(r), between oxygen atoms of water and oxygen (red) and nitrogen atoms (blue) in the PNOMA brush, showing consistent results with previously reported study.^7^; PNOMA: poly(2-(N-oxide-N,N-dimethylamino)ethyl methacrylate)

**Table S1.** Benchmark results from DFT calculations using the B3LYP functional with different levels of theory with Mulliken^3^ and RESP^4^ charge-fitting methods obtained from QChem^5^ and NWChem^6^ software.

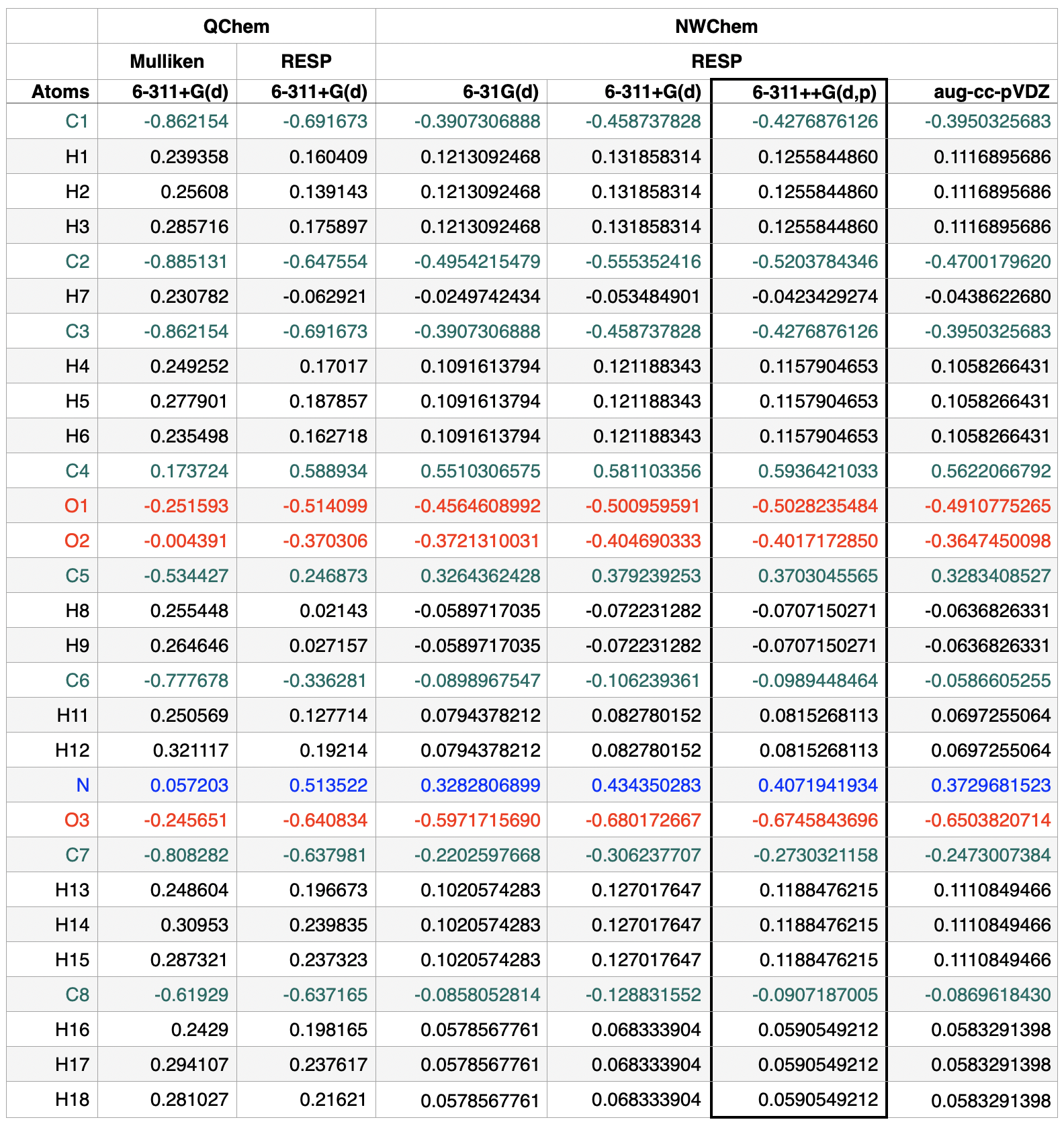

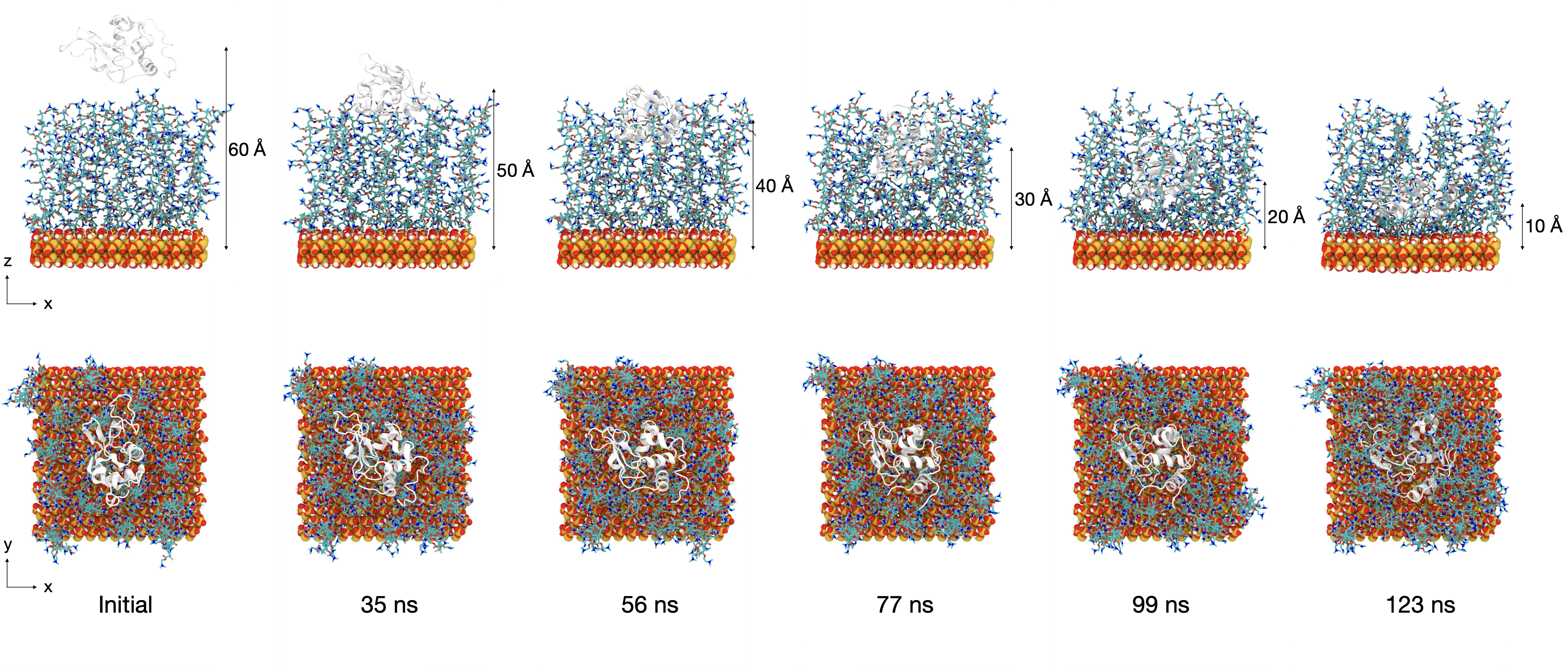

**Figure S2.** Representative snapshots of PDMAEMA brush during the SMD simulation, captured at different distances between the centers of masses (COMs) of lysozyme and the silica substrate, along with corresponding simulation times. The color scheme is the same as **Figure 2**.

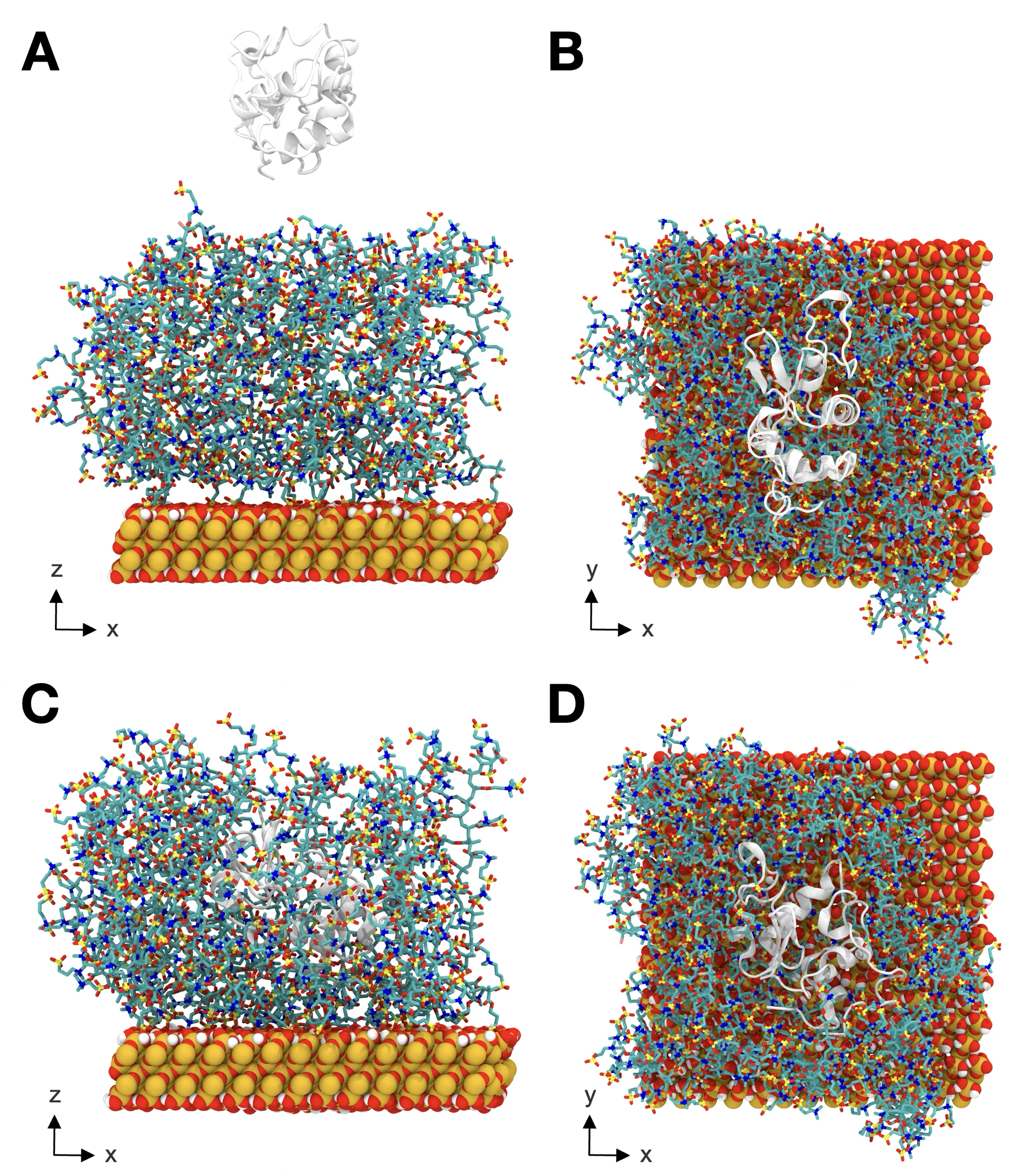

**Figure S3.** (A–D) Representative snapshots of the PSBMA brush at *D*=60 Å and *D*=20 Å, where the distance, *D*, refers to COM separation between lysozyme (light gray) and quartz substrate (orange: Si; red: O; white: H). Side views (A,C) and top views (B,D) are shown. Panels (A,B) correspond to the initial state (*D*=60 Å), while (C,D) correspond to the final state (*D*=20Å).

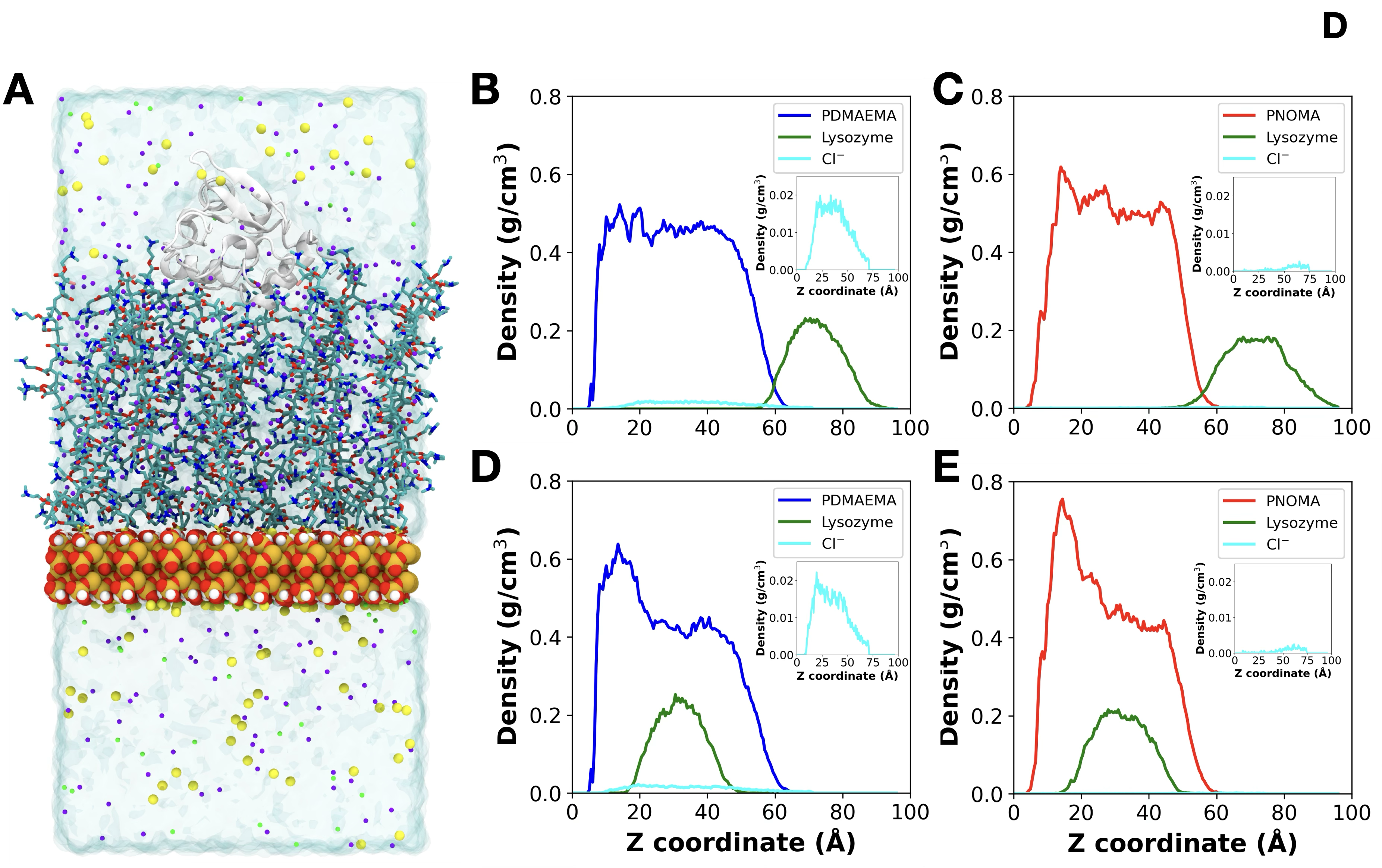

**Figure S4.** (A) A representative snapshot of the PDMAEMA brush system, showing the initial stage of interaction between lysozyme and the brush. The transparent blue box represents water; yellow, green, and purple circles represent Na⁺, K⁺, and Cl⁻ ions, respectively. Other colors follow the scheme used in **Figure 2**. Cl⁻ ions are primarily located within the brush, neutralizing the positively charged PDMAEMA brush. (B–E) Density profiles comparing PDMAEMA (B,D) and PNOMA (C,E) brushes before (B,C) and after (D,E) protein adsorption. The densities of PDMAEMA, PNOMA, and lysozyme are shown in blue, red, and green, respectively. Cl⁻ density (cyan) is enlarged in the insets. The PDMAEMA brush shows a high Cl⁻ density within the brush region (B,D), while the PNOMA brush does not (C,E).
